# Testcrosses are an efficient strategy for identifying *cis* regulatory variation: Bayesian analysis of allele specific expression (BASE)

**DOI:** 10.1101/2020.10.01.322362

**Authors:** Brecca Miller, Alison Morse, Jacqueline E. Borgert, Zihao Liu, Kelsey Sinclair, Gavin Gamble, Fei Zou, Jeremy Newman, Luis León-Novelo, Fabio Marroni, Lauren M. McIntyre

**Affiliations:** Genetics Institute, University of Florida, Gainesville, FL, USA; New York University NY, NY, USA; Department of Molecular Genetics and Microbiology, University of Florida, Gainesville, FL, USA; Department of Biostatistics, University of North Carolina at Chapel Hill, Chapel Hill, NC, USA; Department of Genetics, University of North Carolina at Chapel Hill, Chapel Hill, NC, USA; Department of Pathology, University of Florida, Gainesville, FL, USA; Department of Biostatistics and Data Science, University of Texas Health Science Center at Houston-University of Texas School of Public Health; Department of Agricultural, Food, Environmental and Animal sciences, University of Udine, Udine, Italy

## Abstract

Allelic imbalance (AI) occurs when alleles in a diploid individual are differentially expressed and indicates *cis* acting regulatory variation. What is the distribution of allelic effects in a natural population? Are all alleles the same? Are all alleles distinct? Tests of allelic effect are performed by crossing individuals and comparing expression between alleles directly in the F1. However, a crossing scheme that compares alleles pairwise is a prohibitive cost for more than a handful of alleles as the number of crosses is at least (**n**^**2**^**-n)/2** where **n** is the number of alleles. We show here that a testcross design followed by a hypothesis test of AI between testcrosses can be used to infer differences between non-tester alleles, allowing **n** alleles to be compared with **n** crosses. Using a mouse dataset where both testcrosses and direct comparisons have been performed, we show that ∼75% of the predicted differences between non-tester alleles are validated in a background of ∼10% differences in AI. The testing for AI involves several complex bioinformatics steps. BASE is a complete bioinformatics pipeline that incorporates state-of-the-art error reduction techniques and a flexible Bayesian approach to estimating AI and formally comparing levels of AI between conditions. The modular structure of BASE has been packaged in Galaxy, made available in Nextflow and sbatch. (https://github.com/McIntyre-Lab/BASE_2020). In the mouse data, the direct test identifies more *cis* effects than the testcross. *Cis-by-trans* interactions with *trans*-acting factors on the X contributing to observed *cis* effects in autosomal genes in the direct cross remains a possible explanation for the discrepancy.

## INTRODUCTION

Allele Specific Expression (ASE) is the amount of mRNA each allele transcribes. Allelic imbalance (AI) indicates a difference in the level of expression of transcripts derived from the two alleles of a diploid individual or among alleles in a polyploid (Wittkopp, Haerum and Clark 2004, Boatwright et al. 2018). AI is a result of genetic variation in regulation, both in *cis* (*e*.*g*. promoters, enhancers, and other noncoding sequences), and in *trans* (transcription factors). The interpretation of *cis* and *trans* effects depends upon the experimental design deployed (McIntyre et al. 2006, Graze et al. 2014, Graze et al. 2012, Graze et al. 2009, Fear et al. 2016). Testing for regulatory variation that affect expression in *cis* is conceptually straightforward and involves the comparison of the expression profiles of two alleles with the null hypothesis that the expression profiles are equal. If AI is observed, there is direct evidence of *cis* differences between alleles (Wittkopp et al. 2004). There are also some potential *cis* by *trans* interactions captured in this comparison (Wittkopp et al. 2004, Graze et al. 2014). Comparisons of AI across different physiological or environmental conditions is of increasing interest (Von Korff et al. 2009, Tung et al. 2011, Cubillos et al. 2014, Chen, Nolte and Schlötterer 2015, Buil et al. 2015, Pinter et al. 2015, Fear et al. 2016, Moyerbrailean et al. 2016, Knowles et al. 2017). Formally testing for differences in AI between conditions reveals environmental effects of variation in *cis* regulation (Leon-Novelo et al. 2018) and can also be used to identify parent of origin effects between reciprocal genotypes (Zou et al. 2014).

The direct assessment of AI among **n** alleles would require at least (**n**^**2**^**-n)/2** crosses. This quickly becomes a very large number; for example, directly testing differences between 10 alleles would require 45 crosses. Theoretically it is possible to obtain this information by performing **n** crosses, between one tester inbred line and several non-tester inbred lines to obtain **n** F1 carrying each the same tester allele and different “line” alleles (Figure 1). Testcrosses (or a reference design) are crosses in which two (or more) non-tester alleles are each crossed to a common tester allele. If the non-tester alleles do not differ from the tester, then all three alleles (the two non-tester and the tester) are similar in their effect. If one of the tester/non-tester F1 combinations shows AI but the other does not, this implies that the expression of one of the non-tester allele differs from the other non-tester/tester. What if both alleles differ from the tester? Does this imply these two alleles are similar? A formal test of whether the AI in these two crosses is equal can be used to identify non-tester alleles with divergent *cis* effects. This is an innovative way of estimating allelic effects in a population and predicting alleles likely to differ in *cis*.

**Figure 1:**
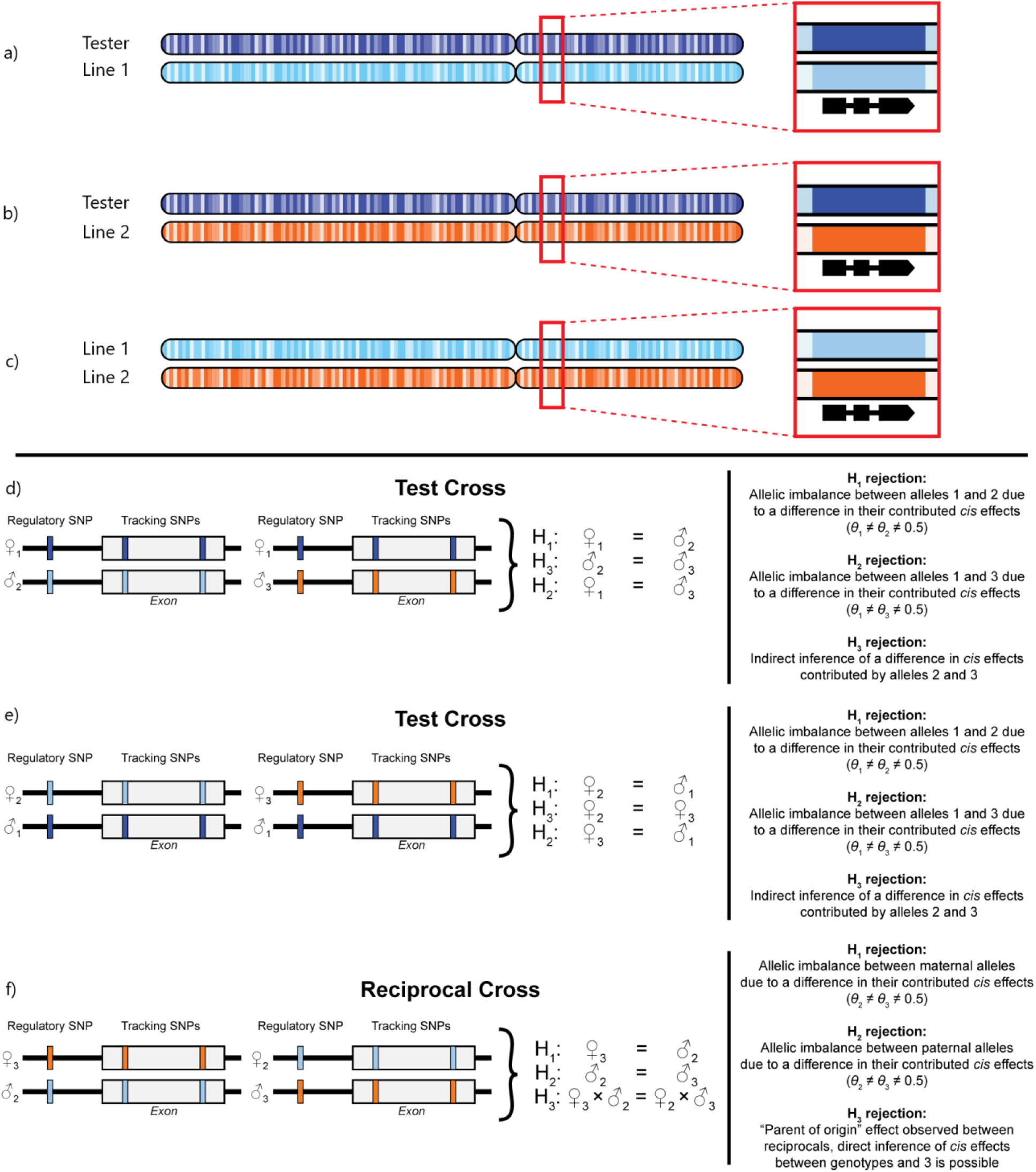
Panels **a** and **b** are F1 progeny from a testcross. The dark blue allele is from the tester parent and is shared between the F1’s while the light blue and orange alleles are the two alleles from the non-tester parents (Line 1 and Line 2, respectively). Panel **c** shows the direct cross between the orange and light blue alleles. Panels **d, e**, and **f** depict genes for which allele specific expression is to be evaluated. The left side shows the genotypes and the right side the corresponding testable hypotheses.

There are many bioinformatics steps needed to get to the point where the expression can be compared across conditions; this causes difficulties in reproducibility and has the potential to discourage researchers without extensive bioinformatics capabilities. Differential mapping of the two alleles on the common reference (Degner et al. 2009) has led to a demonstration that strain-specific reference improves the estimation of AI and lowers type I error (Turro et al. 2011, Skelly et al. 2011, Graze et al. 2012, Satya, Zavaljevski and Reifman 2012, Munger et al. 2014, Leon-Novelo et al. 2014, Fear et al. 2016). This step requires mapping to strain specific references and comparing alignment outputs. Although it should be noted that even with strain-specific mapping, residual bias can persist due to undiscovered structural variants or aspects of sequences that interfere with the mapping algorithms. When DNA can be used as a control, all of these biases are accounted for (Wittkopp et al. 2004, Graze et al. 2009, Graze et al. 2012) but some residual bias may persist for example when gene families are considered. Bias due to sequence similarity across the genome and/or mapping algorithm can be estimated using simulation (Degner et al. 2009, Stevenson, Coolon and Wittkopp 2013, Leon-Novelo et al. 2014) these estimates can be accounted for in the model reducing type I error (Graze et al. 2012). Other bioinformatic challenges include tracking overall expression and non-specific read mapping, as they both affect the power for detection of AI (Fear et al. 2016, Leon-Novelo et al. 2018).

Here, we present BASE, a series of clearly elucidated bioinformatic steps modularized and well documented in a robust high performing computing (HPC) pipeline. BASE requires only reads in FASTQ file format, a reference genome, genotype specific VCF file for variant calling, and a series of design files for as input. The tools within the pipeline are modular to provide a clear and reproducible analysis workflow. The template provided makes for transparent substitution and easy execution. Workflow templates have been constructed for SLURM schedulers, for Nextflow (Di Tommaso et al. 2017), and for Galaxy (Goecks et al. 2010). Galaxy BASE consists of individual tools, associated workflows and a detailed conda environment that is available as a PyPi package for integration into local Galaxy installs or servers. Individual BASE tools and workflows are also available for installation from the Galaxy ToolShed.

While this paper focuses on the use of testcrosses for comparison, BASE is entirely general. The first design file specifies individual crosses and the corresponding genomes. Any crossing design can be accommodated and the pipeline will then automatically assemble the relevant genomes and count alleles mapping to each parental genome. Further the unit for which allele specific reads are to be counted is completely general as the process expects a labeled BED file. This is used throughout the process to identify results for each row of the user provided BED file. Pairwise comparisons are specified in a separate design file and the only limitation is the requirement that the identifiers on the genomic regions match. For example, two tissues/environments from the same genotype, two sexes, reciprocal crosses and the testcrosses presented here can all be analyzed with the same process. Our Bayesian model allowing simultaneous estimation of AI in two conditions and direct estimation of the difference in AI between conditions (Leon-Novelo et al. 2018), has been recoded in Stan (Carpenter et al. 2017) making the code more robust. Finally, as with the other components a custom model for AI in a particular circumstance can easily be deployed (Zou et al. 2014) using modules in BASE to count allele specific and non-specific reads.

## METHODS

### BASE Modules

BASE consists of four main modules: *Genotype Specific References, Alignment and SAM Compare, Prior Calculation*, and *Bayesian Model* (Figure 2). Flexibility, generality and process checking are present in each of the four modules that make up the BASE pipeline. Workflows for each module are coded in sbatch, Nextflow and Galaxy, and all scripts are available on Github (https://github.com/McIntyre-Lab/BASE_2020). The Galaxy package has a detailed User Guide (Supplementary file) that also describes the structure of the workflow and all input and output requirements for each of the individual scripts.

**Figure 2:**
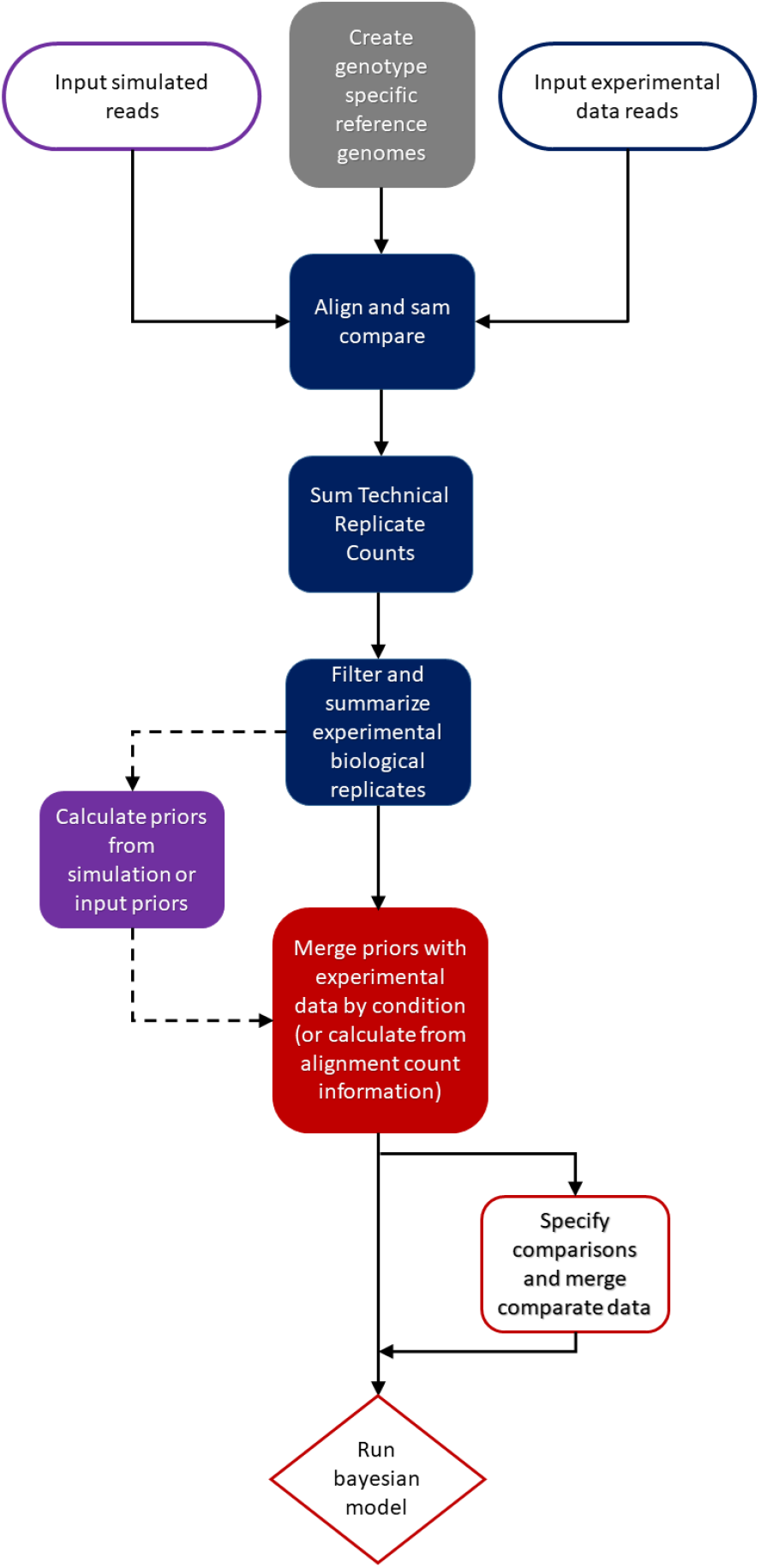
A flowchart of the Bayesian analysis of allele imbalance (BASE) process for testing for allele imbalance. BASE is divided into four distinct modules. The modular structure allows the user to follow each step, and as new tools become available to substitute components. The design flexibility allows users to specify their own comparisons allowing multiple different genetic designs to be accommodated in a straightforward manner. on the user’s preferred method of creating genome-specific reference genomes. Each module has a clearly defined set of input an output files and the user can elect to use any individual module or the entire pipeline. Design files structured formats are used to automating the bioinformatic steps while enabling complete flexibility in the genetic design. The details are described in the user guide (supplementary material) and the process is implemented in SBATCH, Nexflow and Galaxy (https://github.com/McIntyre-Lab/BASE_2020)

The *Genotype Specific References* module (Supplementary Figure S1) requires as input the reference genome as FASTA file, and genotype specific SNP variants as VCF files, and returns as output a set of genotype specific references obtained incorporating SNP variants into the reference genome. Genotype specific reference reduce mapping bias that occurs when a common reference is used (Turro et al. 2011, Skelly et al. 2011, Graze et al. 2012, Satya et al. 2012, Munger et al. 2014, Leon-Novelo et al. 2014, Fear et al. 2016). Input VCF files can be generated using the GATK (DePristo et al. 2011), and index input genotype VCF files. BWA index (Li and Durbin 2010) is used to create index files needed for downstream alignment.

The *Alignment and SAM Compare* module (Supplementary Figure S2) quantifies alignment counts for each input file for each of the two genotype specific genomes of the parents of the F1, compares the alignment files in SAM format, and outputs count tables of reads aligned to each parental genome. In this module, input experimental reads for each F1 sample are aligned to each of their updated parental reference genomes with BWA-MEM (Li 2013). Alignment output counts are compared and reads are designated as mapping to one of the parental genomes (or to both when reads mapping equally well to the two genomes). Technical replicates are summed together, and coverage metrics are calculated. The data are then flagged for low coverage (user defined).

The *Bayesian model* requires an estimate of a prior to specify the probability that a read generated from the gene in genome 1 maps better to genome 1 than to genome 2 or is mapping equally well on both genomes for each condition. In this implementation, the prior is specified for each gene/cross and each condition separately allowing for maximum flexibility. For each gene/cross/condition, *q1* is the prior probability that a read originating from genome 1 maps better to genome 1 and *q2* is the prior probability that a read originated from genome 2 maps better to genome 2, with the following constraint: 0<*q1,q2*<1. The *Prior Calculation* module (Supplementary Figure S3) estimates priors from DNA data, the RNA data, or simulated data. This module receives count tables for read counts for both informative (mapping to a particular parent) and uninformative reads as input and uses them to estimate a prior probability distribution for each given feature. Priors can also derive independently from this tool and supplied directly to the model. Priors provide information on differences in mapping on the two genomes in absence of AI (Graze et al. 2012, Fear et al. 2016, Leon-Novelo et al. 2018).

The *Bayesian Model* module (Supplementary Figure S4) requires as input a design file to identify comparisons to be performed, alignment count tables and priors, either from the previous module or as direct input. The Bayesian model itself is the STAN (Carpenter et al. 2017) implementation of a previously published model for the detection of AI in one or between multiple conditions (Leon-Novelo et al. 2018) with an extension allowing more flexibility in the prior. Specifically, the previous implementation assumed that the prior was the same for both conditions, here this assumption was relaxed and the priors are now independently specified between conditions. The model was developed and tested without that assumption and the restriction was only in the previous R-code for model implementation. Model output files include values for estimates of levels of AI and their 95% central credible intervals. Output also includes the Bayesian measure of evidence against allelic balance (Thulin 2014, de Bragança Pereira and Stern 1999), defined as the smallest number *ev* such that the 1-*ev* central credible interval for estimate of AI does not contain the null value indicating allelic balance. Values of *ev* can be used in a decision theory context to make decisions about rejecting the null hypothesis.

### Workflow Deployment Platforms and Protocol

To facilitate its use, BASE is available for Galaxy, as a workflow on the Nextflow platform, and with an example set of sbatch scripts for SLURM schedulers on GitHub (https://github.com/McIntyre-Lab/BASE_2020). Source code is available under the MIT license. BASE has a modular structure; flowcharts of the individual modules give details about the logic (Supplementary Figures 1-4) and are described in detail in the User Guide (Supplementary Material). Names are consistent with the flow charts and standard naming conventions for sbatch and Nextflow are used.

Galaxy is designed to be deployed in a web browser, providing a user-friendly platform that can be configured to run on global servers in large universities or on the local individual machines (Goecks et al. 2010). Reproducibility in Galaxy is accomplished via histories and the creation of workflows that can be shared among collaborators. Nextflow (Di Tommaso et al. 2017) is a portable, parallelizable and reproducible framework. It defines data workflows that can be executed on diverse portable batch system schedulers such as SGE, SLURM or Cloud platforms. Nextflow pipelines consist of a configuration file and a series of processes that define the major steps in the pipeline. Processes are independently executed from one another and the platform supports a variety of languages such as Bash, Python, R, etc. Processes are connected through input or output channels, allowing data to be passed through the pipeline. Nextflow also provides an automatic caching mechanism for identifying and skipping successfully completed tasks and using previously cached results for downstream tasks.

The third option that is provided for users is through several templates for sbatch scripts; they can be modified as needed with the input and output file names and submitted to a SLURM scheduler or used as templates for deployment using a different scheduler.

### Testcrosses can be used to efficiently estimate cis effects

To demonstrate that the testcross can be used to predict allelic differences between non tester alleles we analyzed a publicly available dataset of male and female F1 *Mus musculus* brain RNA-seq from three different inbred mouse strains, CAST/EiJ, PWK/PhJ, and WSB/EiJ (Zou et al. 2014, Crowley et al. 2015). The authors provided two alternative notations for the inbred strains: CAST/EiJ = F, PWK/PhJ = G, and WSB/EiJ = H, or alternatively CAST/EiJ = CAST, PWK/PhJ = PWK, and WSB/EiJ= WSB. In the present work, and similar to what has been done previously, we denote each F1 sample by its maternal strain X paternal strain. For example, a CAST X WSB mouse is an offspring of a CAST/EiJ female that is mated with a WSB/EiJ male. This same F1 sample can be denoted as FH (short for F X H) in which an F male is mated with an H female. This shorter notation is used to label some tests/figures and the model results (Supplementary file SF01). For the original study the main hypothesis of interest was a test of the null AI_FG_ =AI_GF_, AI_GH_ =AI_HG_, AI_FH_ =AI_HF_, *i*.*e*. the absence of a parent-of-origin effect. We repeated the analysis with BASE, using the same read counts as the original work, and male and female offspring of each cross were analyzed separately as a series of pairwise analyses using the Bayesian model (Leon-Novelo et al. 2018) and not jointly (Zou et al. 2014).

Our main interest was to assess whether the theoretical prediction that the testcross can be used to compare different non-tester alleles was examined. In the mouse data, the offspring of the crosses CAST X PWK and CAST X WSB, represent testcrosses with CAST as the tester allele. Using the flexible design in BASE, we estimated AI and tested whether the AI was significant between alleles CAST and PWK in the offspring of CAST X PWK (H1 in Figure1d); alleles CAST and WSB in the offspring of CAST X WSB (H2 in Figure 1d), and the difference in AI between the two offspring (H3 in Figure 1d). The null hypothesis H3 is that the expression of PWK in CAST X PWK is the same as the expression of WSB in CAST X WSB. Since CAST is common to both crosses (and the maternal allele, in this case), testing H3 is a test between PWK and WSB. We also compared the test crosses PWK X CAST and WSB X CAST in this case CAST is still the tester allele but here it is inherited paternally. The other two testcross combinations were tested in the same manner. We focus only on the autosomal effects and do not consider the X, Y or mitochondrial loci.

To verify the predictions made in the testcross, the direct crosses were used (Figure 1d). For example to verify differences in the PWK and WSB alleles predicted by the CAST X PWK and CAST X WSB cross we examined the PWK X WSB and WSB X PWK crosses. Since the alleles in the testcrosses are compared with the same parental inheritance and the alleles in the direct test have different parental inheritance we do not expect that 100% of the predictions will be realized, as those predictions do not take into account parent of origin effects. The Spearman coefficient of correlation was used to compare the estimates for AI estimated from BASE with the estimates of AI used in the original work (Zou et al. 2014, Crowley et al. 2015) for the 95 imprinted genes and all data were analyzed using the same counts from the original analysis.

## RESULTS

As expected, estimates of allelic effect obtained with BASE were concordant with the published estimates of AI from the 95 imprinted genes (Zou et al. 2014, Crowley et al. 2015). Figure 3 shows, for the reciprocal CAST X PWK and PWK X CAST crosses, the estimated AI as the ratio of the paternal to the maternal allele. All the correlations coefficients are greater than 0.96, even for reciprocal crosses. The correlation for the same cross in the present study and published results is always 1. Using the analysis in BASE we report the *cis* effects as estimated by tests of AI for each cross. The estimates range from 5.39% to 11.74% of genes showing AI for each cross (Figure 4) and an overall estimate of 3,629 (26%) loci of the 14, 058 that were tested for all crosses in both sexes. As these data have been analyzed and reported on elsewhere (Zou et al. 2014, Crowley et al. 2015), we will focus on the testcross results.

**Figure 3:**
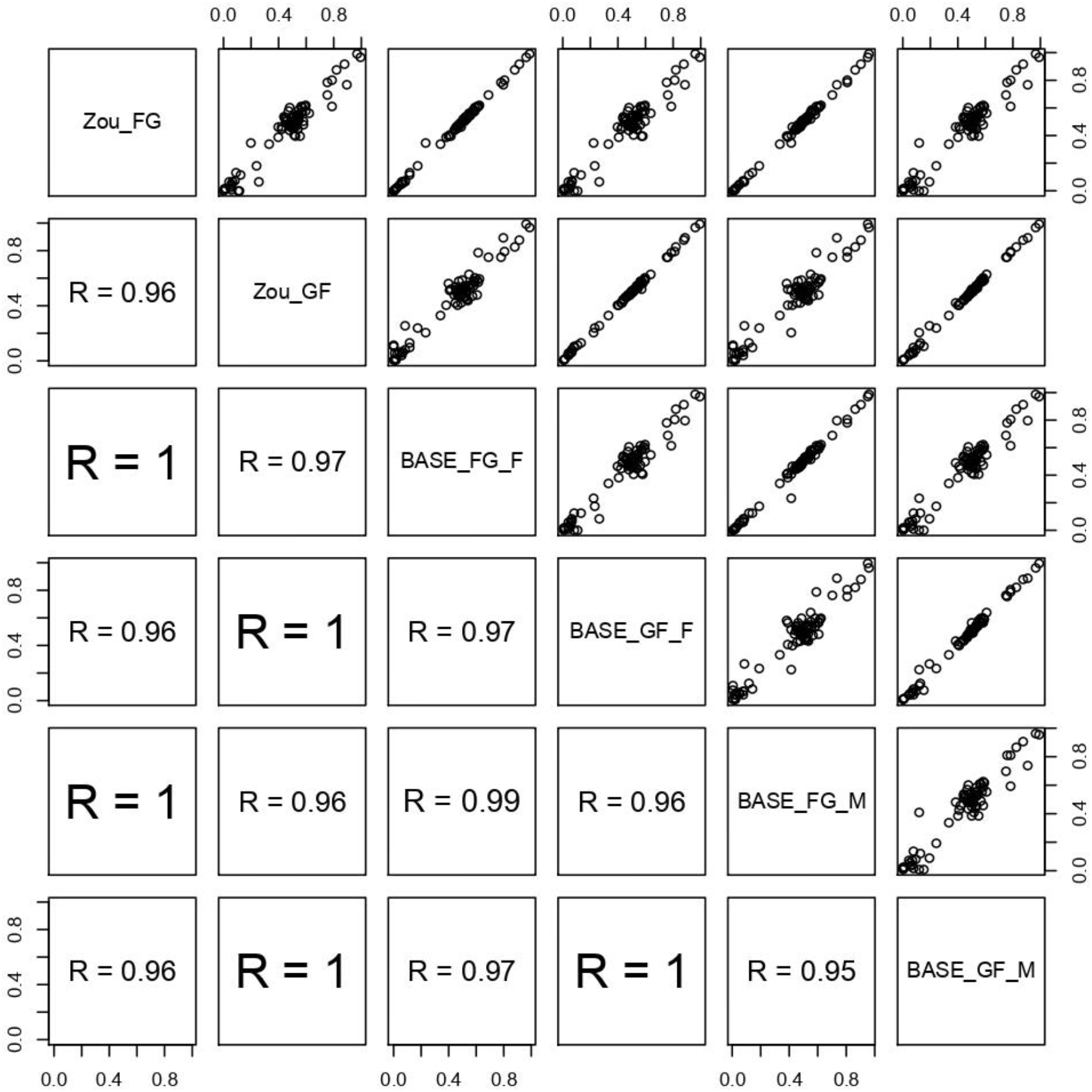
The correlation in estimates for 95 imprinted genes between the original data analysis (Zou) and the Bayesian model implemented in the present work (BASE). In the upper triangle are the correlation plots with the BASE estimates on the X axis and the estimates from the original data analysis (Zou et. al. 2014) on the Y axis. In the lower triangle are Spearman’s correlation coefficients. **FG** = CAST X PWK, **GF** = PWK X CAST, **F** = Females, **M** = Males

**Figure 4:**
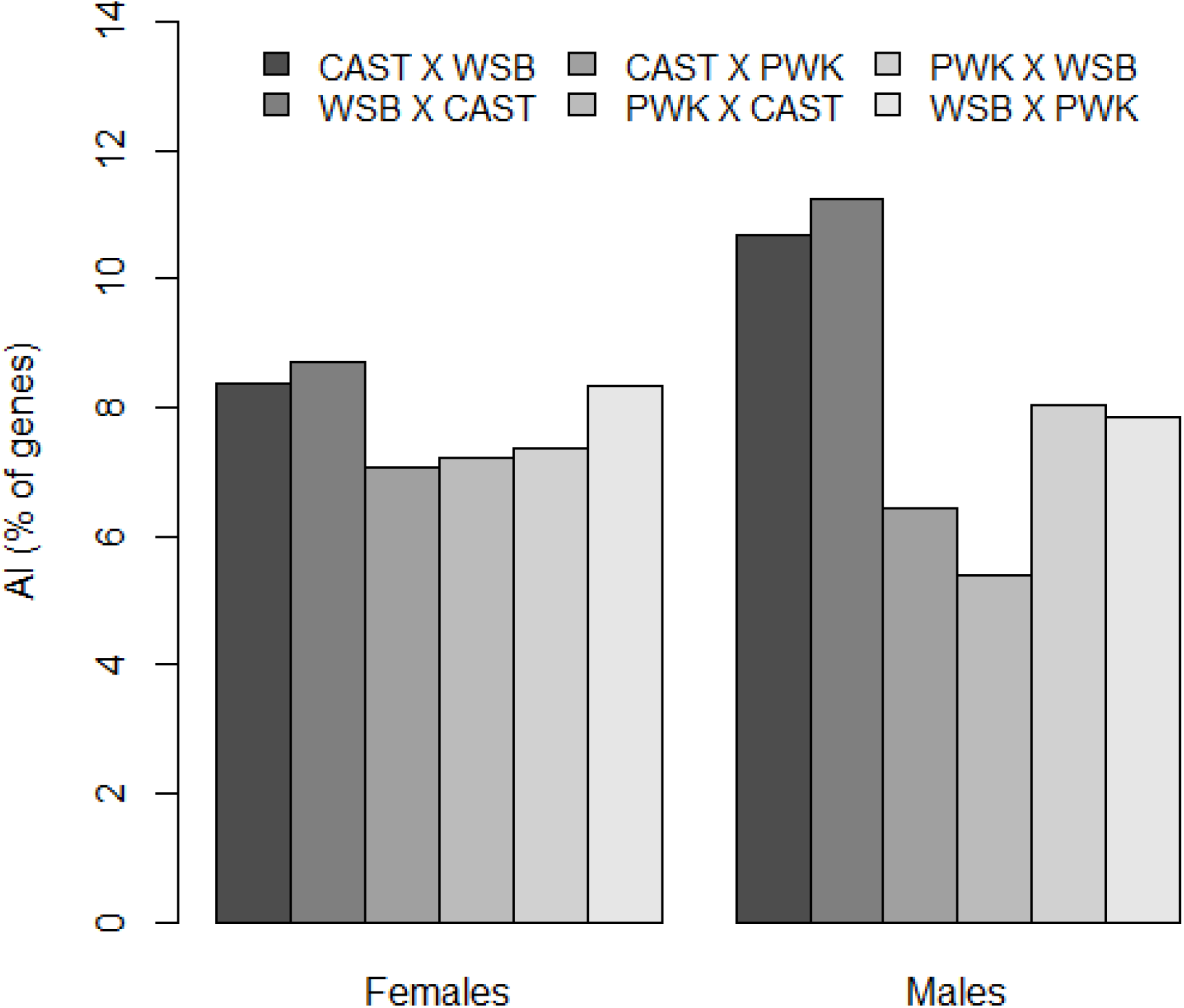
Percentage of genes showing AI in individual crosses.

The testcross predicts ∼1,200 loci with *cis* effects in PWK X WSB, ∼900 loci with *cis* effects in CAST X PWK and ∼1000 in CAST X WSB, accounting for 2% to 5% of the tested genes (Figure 5). The frequency of cis effects was similar regardless of the parent of origin. Using BASE, we directly tested the null that the AI between the sexes was equal for each of the 6 F1 crosses. The detection of cis effects was similar for male and female offspring with the exception of the CAST x WSB cross where AI was 2x more likely to be identified in male offspring compared to female offspring (p<0.0001 McNemar’s test).

**Figure 5:**
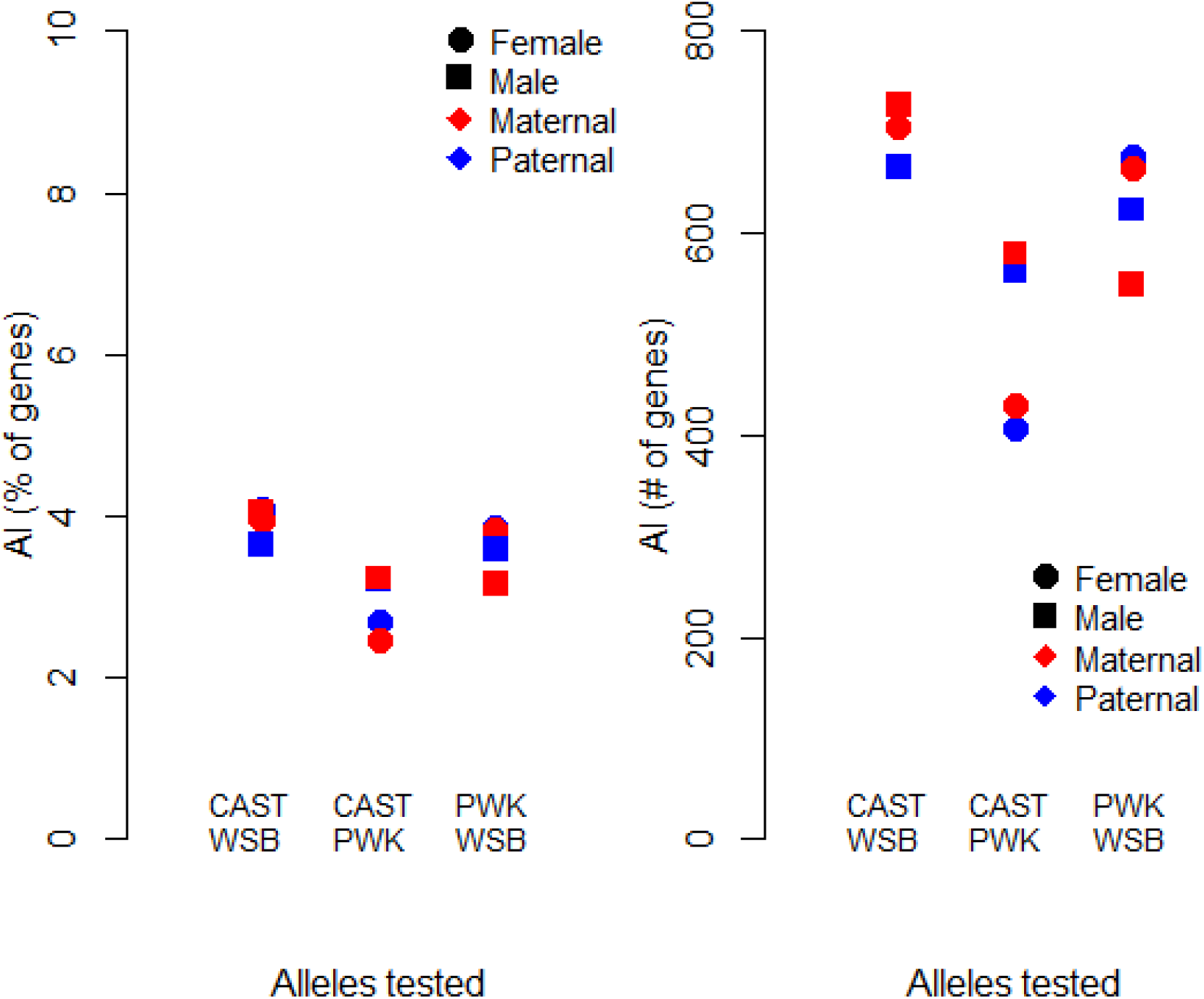
Allele specific expression between pairs of alleles, tested via the testcross approach. The X axis represents the two alleles being compared. CAST and WSB are compared by testing H3 in the two crosses CAST X PWK and WSB X PWK (to test Maternal contribution, red) or PWK X CAST and PWK X WSB (to test Paternal contribution, blue). Results for female offspring are shown as circles, and fro male offspring are shown as squares. On the left, we report the percentage of genes showing AI, and on the right we show the absolute values.

Remarkably, more than 70% of *cis* effects predicted by the testcross as differences between non-tester alleles (*i*.*e*. rejection of the null hypothesis H_3_) are validated in the direct comparison (Figure 6) for all comparisons except the CAST X WSB male which still has a validation rate of 46%, or approximately 3 times higher than expected based on the frequency of *cis* effects in the direct cross. We also examined whether the presence of a parent of origin effect changed the validation rate and the validation rate increased when parent of origin effects were present (data not shown).

**Figure 6:**
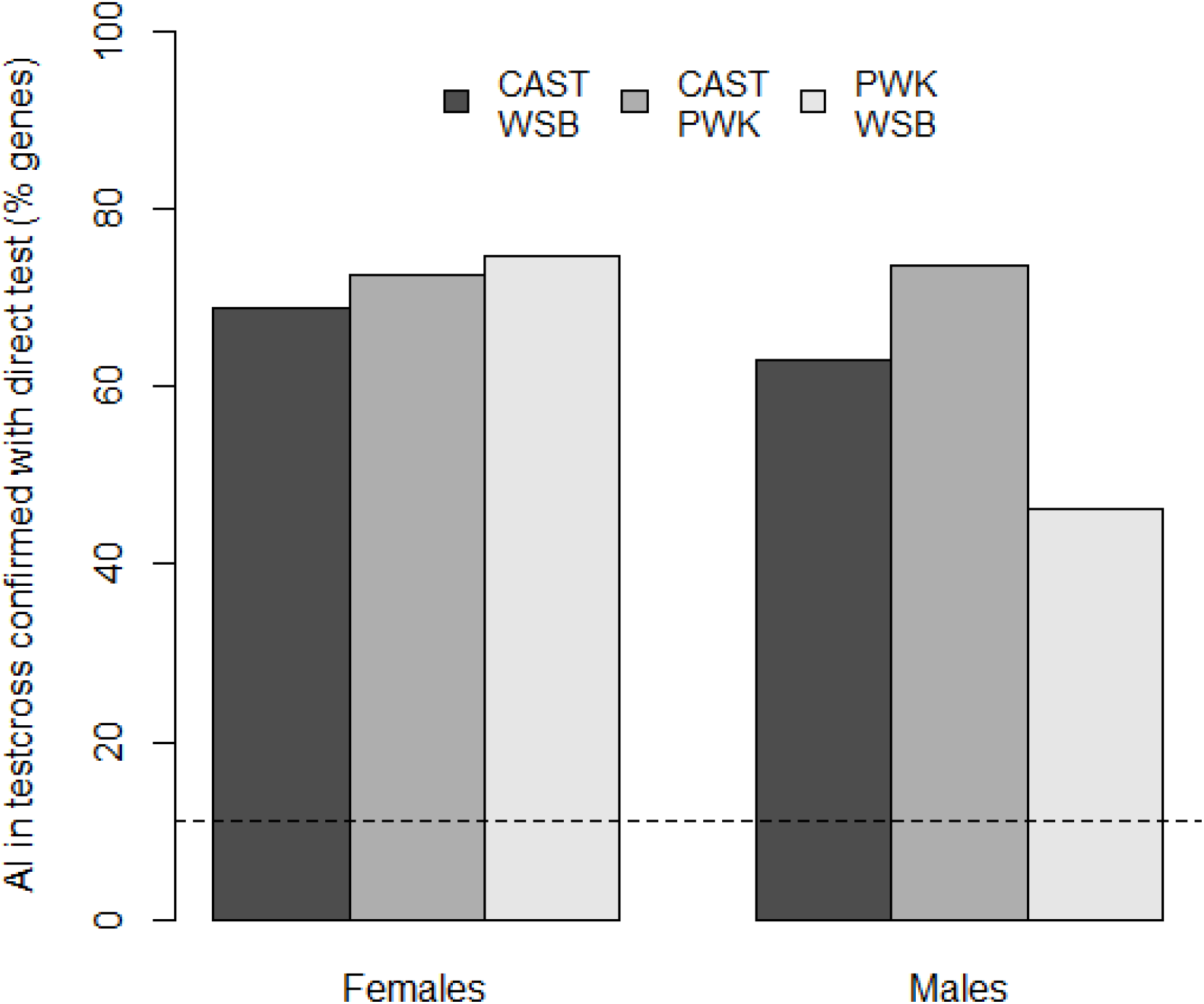
Percentage of *cis* effects detected in the testcrosses, validated in the direct cross in female and male offspring. Proportion expected by chance indicated by the horizontal line estimated as the maximum marginal frequency among the direct crosses. (11.2%)

## DISCUSSION

BASE has been used here to test for *cis* effects in individual crosses, differences in *cis* effects between males and females of a single cross, differences in *cis* between testcrosses, and parent of origin/cis-trans interactions in reciprocal crosses. The BASE framework lays out each step to testing AI transparently, and enables researchers to perform analysis using their preferred approach (galaxy, Nextflow, or SBATCH queuing). We provide a set of well-documented python scripts organized into modules and available with examples as SLURM bash jobs, using Nextflow or Galaxy. BASE its designed in a modular fashion. Users can rely on the whole pipeline of analysis, select a specific step, or replace a specific step. The addition of usable accessible code should make these more complex models and bioinformatics steps more accessible to the community.

Using a testcross design is an efficient way of identifying *cis* effects. Instead of a joint pairwise **n**^**2**^**-n** experiment it is possible to plan an experiment of size **n**, where **n** is the number of alleles tested in one sex or paternal/maternal effects. There are some subtleties worth considering when thinking about *cis* effects in organisms with a heterogametic sex system. The testcross will compare non-tester alleles inherited from either the maternal or paternal parent. The direct comparison, in contrast, will always compare the maternally inherited allele to the paternally inherited allele. When there is no parent of origin effect, and no interactions between the heterogametic sex chromosome or cytoplasmic factors and the autosomes, this should result in the same loci being identified. However, a parent of origin effect, *trans* acting factors from the heterogametic sex chromosome and cytoplasmic factors complicates the interpretation of the direct comparison, particularly in males. In the comparison between testcrosses there is potential for reduced power, for example if the coverage between the two testcrosses is unequal.

The mouse data show that the use of testcrosses to compare non-tester alleles identifies loci that are validated by the direct tests in the vast majority of cases. Test crosses that survey both maternal and paternal inheritance and identify common loci are validated at an even higher rate. Loci identified in direct crosses are identified in test crosses but there are many more loci identified in the direct cross. This, may be a false negative result for the testcross approach, due to lower power. However, it may well be that there is a *cis-trans* interaction between the heterogametic sex chromosome and the autosomal genes, and/or a parent of origin effect in the direct cross.

The parent of origin effect can be tested by comparing AI between reciprocal crosses. Here the original study focused on the parent of origin effect, and so we can examine whether this effect explains some of the differences in the *cis* effects estimated in the testcross compared to identified in the direct cross. For 60%-75% of the loci with a parent of origin effect there was evidence for a *cis* effect in the direct cross as well, with the exception of the CAST X WSB male where 46% of the parent of origin effects had corresponding *cis* effects. This is logical as at least one of the reciprocal crosses must have a relatively large estimate of AI in order to detect the parent of origin effect. It is not unexpected that the validation rate of loci with *cis* effects as estimated from the testcross are even more likely to be validated than those without evidence for a parent of origin effect.

*Trans* acting factors and likely *cis-trans* interactions may explain why more *cis* effects are detected in the reciprocal cross than in the testcross. In the original analysis several hundred genes were differentially expressed between strains on the X chromosome. X-linked loci have been shown to regulate autosomal gene expression such as the non-coding loci *Firre* and *Dxz4* in both male and female mice (Hacisuleyman et al. 2014, Andergassen et al. 2019). Inheritance of the X in females from different parents, and in males the X is different in the two reciprocals. This effect was described in the original papers (Zou et al. 2014, Crowley et al. 2015).

The large differences in CAST X WSB male are not as surprising as at first glance. This is a cross between genetically distant lines (Zou et al. 2014) and may reflect divergence in gene expression between the sexes in these incipient species due to sex antagonism. Alternatively, the data may represent a different potential biological explanation for the divergence between the male and female results for this cross.

The testcross approach is a useful strategy to maximize allele comparison while minimizing sequencing efforts. Testcrosses will not detect either parent of origin or *cis-trans* interactions since the comparison between alleles is from a shared maternal/paternal inheritance. The reciprocal effect is large in these data indicating that either parent of origin and/or *cis-trans* interactions are important in these data, consistent with the original data analysis (Crowley et al. 2015).The existence of X signaling effects lends support to this hypothesis. Other work has also implicated *trans*-acting factors from the X influencing allele specific expression on the autosomes (Graze et al. 2014). The efficacy of the testcross is clear from these data, also clear is the presence of *cis-by-trans* effects from the X, mitochondrial or Y chromosome influencing expression variation in autosomal genes in the mouse.

## ACKNOWLEDGEMENTS

Olexander Moskalenko for critical help on Galaxy infrastructure. Alexis Earley contributed to the Galaxy pipeline, and Adalena Nanni for helpful edits. NIH NIGMS GM128193. HiPerGator High Performance Super Computer at the University of Florida.

## DATA AVAILABLILITY

All scripts and documentation are available at https://github.com/McIntyre-Lab/BASE_2020. Results from all analyses conducted are included as a supplementary file. The mouse data are available from prior publications (Zou et al. 2014, Crowley et al. 2015).

