## Supplementary_Figures for "Testcrosses are an efficient strategy for identifying *cis* regulatory variation: Bayesian analysis of allele specific expression (BASE)"

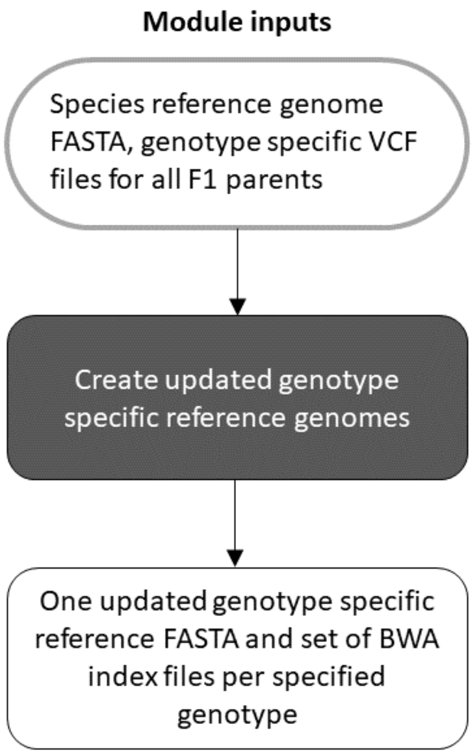


**Figure S1:** Workflow of the Genotype Specific Reference Module.

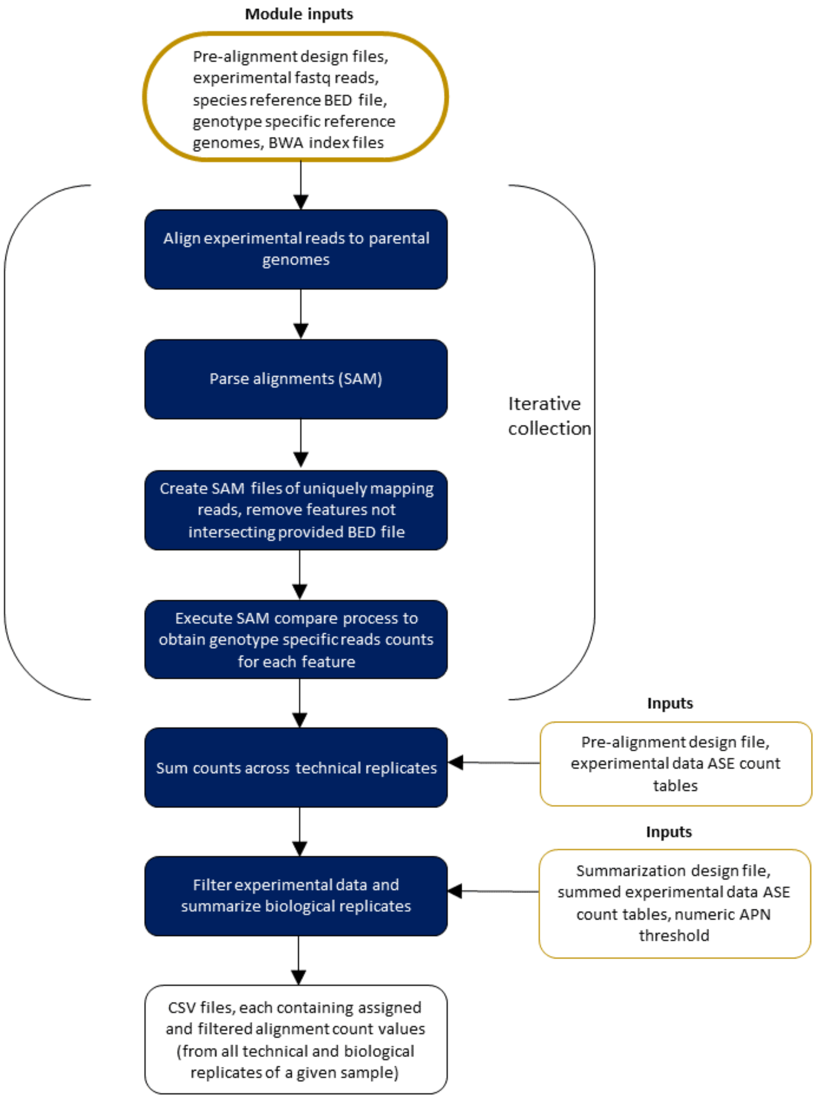


**Figure S2:** Alignment and SAM compare Module.

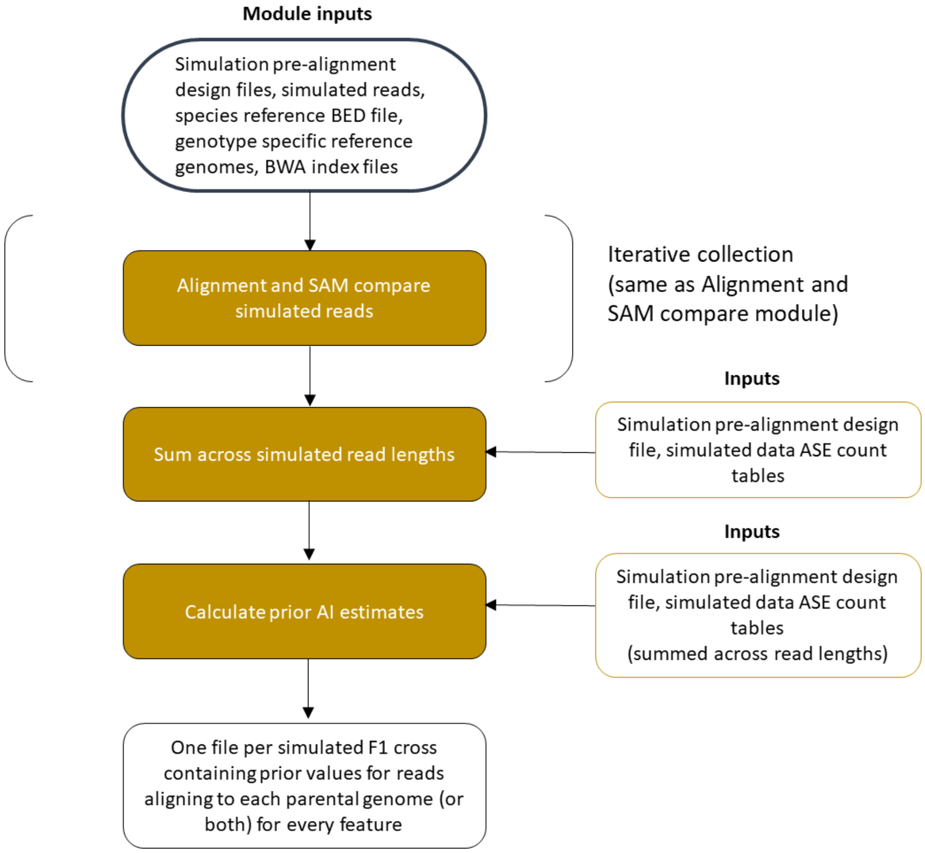


**Figure S3:** Prior calculation module.

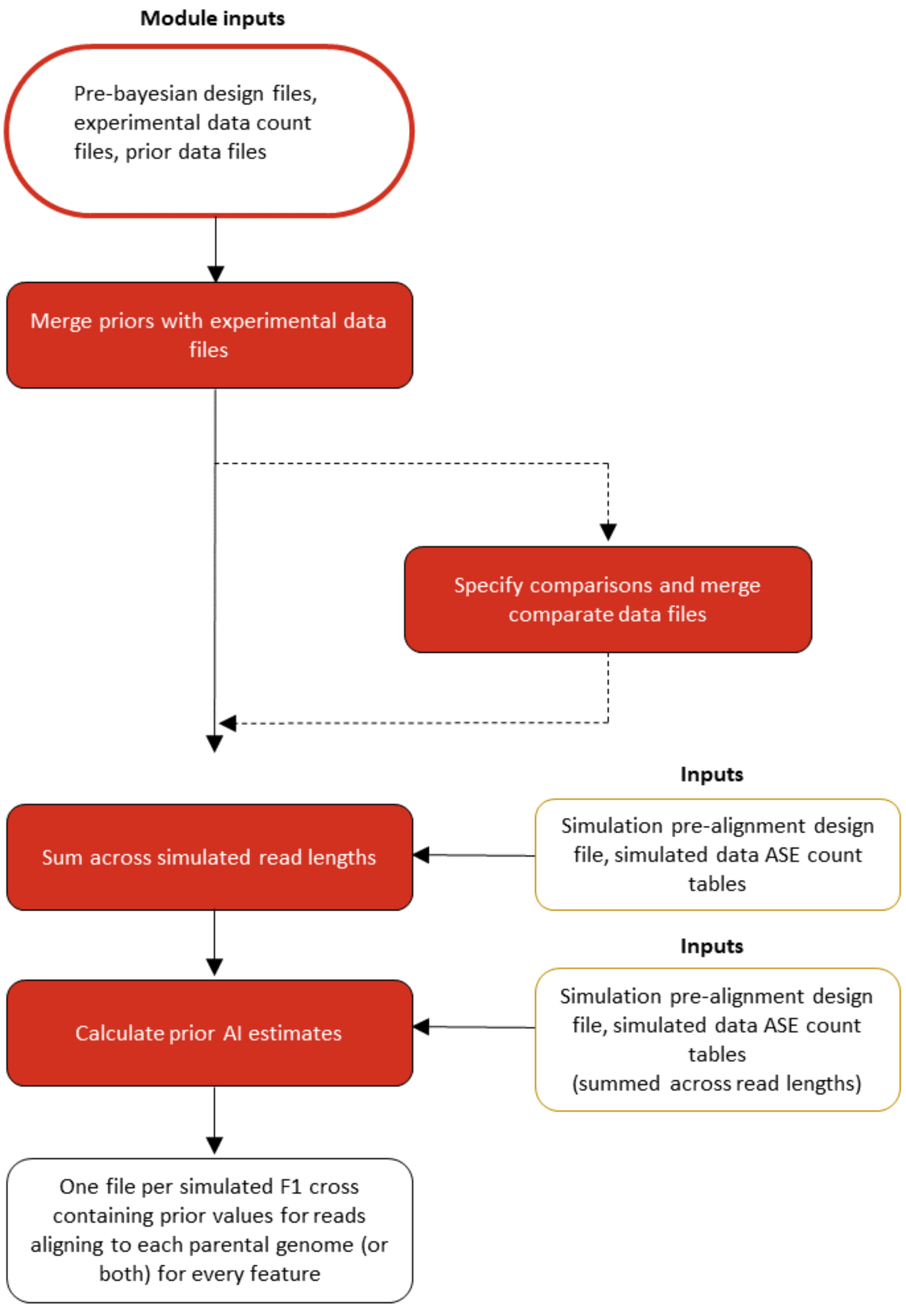


**Figure S4:** Bayesian Model module.
